# Enhancing Transcriptional Data Reliability in Fish Oogenesis Using cDNA-Based Normalization

**DOI:** 10.64898/2026.03.26.714387

**Authors:** Iratxe Rojo-Bartolomé, Jone Ibañez, Ibon Cancio, Maren Ortiz-Zarragoitia, Eider Bilbao

## Abstract

Transcriptomic analyses are widely used to elucidate the molecular mechanisms driving gametogenesis and reproduction in fish, yet their accuracy depends heavily on appropriate normalization of gene expression data. Conventional approaches that rely on single or multiple reference genes are problematic during teleost oogenesis, as profound structural and physiological remodeling of the ovary challenges the assumption that commonly used reference transcripts remain stable. In this study, we assessed by qPCR the transcriptional variability of four widely used reference genes (*actb*, *ef-1α*, *gapdh*, and *18S rRNA*) throughout the oogenic cycle of the thicklip grey mullet (*Chelon labrosus*), using geNorm and NormFinder analyses, and we additionally evaluated total cDNA concentration as an alternative normalization factor. To examine the performance and interpretive consequences of each normalization strategy, we compared expression patterns of key steroidogenic genes (*star*, *cyp19a1a*, and *cyp11b*) normalized by individual reference genes, combinations of reference genes, or total cDNA concentration. All evaluated reference genes displayed notable transcriptional variability across oogenesis, confirming their limited suitability as sole internal controls. In contrast, normalization approaches integrating multiple reference genes and/or total cDNA concentration consistently provided greater stability and more reliable biological interpretation. These results support a refined and more robust normalization framework for transcriptional analyses in fish ovaries, particularly during stages of extensive tissue remodeling. Our findings demonstrate cDNA-based normalization is straightforward, rapid, and easy to implement across laboratories, providing a practical alternative for achieving accurate, reproducible transcript quantification in fish ovary studies.

## INTRODUCTION

Quantitative real-time PCR (qPCR) has become a widely used and essential technique for investigating cellular responses to diverse physiological and environmental stimuli. As a highly sensitive and rapid method for quantifying mRNA levels, qPCR enables precise measurement of gene expression (Taylor et al., 2019). However, this same sensitivity requires strict control of experimental conditions to ensure the accuracy, reproducibility, and comparability of results across samples and studies. To promote methodological rigor, the MIQE (Minimum Information for Publication of Quantitative Real-Time PCR Experiments) guidelines were introduced, providing detailed recommendations for experimental design, data acquisition, and reporting standards (Bustin et al., 2009). These guidelines emphasize, among other aspects, the importance of selecting appropriate normalization strategies to obtain biologically meaningful gene expression data. Despite these efforts, the choice of an optimal normalization method remains challenging and continues to be a matter of debate, as reference gene stability can vary across tissues, developmental stages, and experimental conditions (Dheda et al., 2005; De Santis et al., 2011; Song et al., 2022). More recently, the updated MIQE 2.0 guidelines (Bustin et al., 2025) have expanded the original framework to reflect technological advances in qPCR reagents, instrumentation, and clinical applications. These updated recommendations further highlight the need for enhanced reproducibility and transparency in data interpretation—issues that remain central to ongoing discussions within the scientific community (Bustin et al., 2025).

qPCR data are commonly corrected using three principal approaches based on the cycle threshold (CT), defined as the amplification cycle at which the fluorescence signal surpasses the baseline and which is inversely proportional to the initial amount of mRNA or DNA. The first approach, the ΔCT method (e.g., Kroupová et al., 2011), normalizes target gene expression to the initial quantity of total RNA or mRNA. The second and most widely applied approach is the ΔΔCT method (Pfaffl et al., 2001; Schmittgen & Livak, 2008), which normalizes target gene transcription relative to that of an internal reference or housekeeping gene. The third method also builds on the ΔΔCT framework, but instead of using a single reference gene, it employs the geometric mean of multiple reference genes to achieve a more stable normalization factor (Vandesompele et al., 2002; Bustin et al., 2025). Each of these approaches presents specific advantages and limitations. The ΔCT method, which relies on RNA quantification, is the simplest and least expensive; however, it has important constraints. Spectrophotometric measurements at 260 nm, commonly used to quantify RNA, have low sensitivity and cannot distinguish nucleic acids from contaminants or residual genomic DNA introduced during extraction (Hashimoto et al., 2004; Lundby et al., 2005), particularly when DNase treatment is omitted. Although posterior fluorescence-based RNA quantification methods provide improved sensitivity, they do not account for variability stemming from inefficiencies in cDNA synthesis (reverse transcription, RT) or qPCR amplification, both of which should be minimized (Bustin et al., 2025). It is well established that the RT step is a major source of experimental error (Ståhlberg et al., 2004), and because this step follows RNA extraction, any normalization strategy relying solely on RNA quantity is inherently less reliable (Lundby et al., 2005).

The use of internal reference genes shows an important advantage, as it takes into account errors made during the experimental procedure. Thus, over the years the use of internal reference genes has been the most widely used method for the correction of transcription levels, being even recommended in the MIQE guidelines. However, this method also shows its limitations. On the one hand, internal reference genes must be chosen with great care and cover at least three main conditions: (1) the gene(s) must be constantly transcribed in different cells and (2) under different experimental conditions, and (3) its/their transcription levels must be similar to the transcription levels of each target gene (Zhu et al., 2008). Therefore, there is no universal reference gene, and validation sessions must be performed with each new experiment, species, or organ / cell, or condition (Gutierrez et al., 2008; Hugget et al., 2005). In fact, the MIQE guidelines do not recommend using unvalidated genes and in addition, it favors the use of more than one reference gene, allowing the use of a single gene only in certain specific cases (Bustin et al., 2009; 2025). All this greatly increases the cost of the experiment (Libus and Štorchová, 2006) but as a result, many research papers using a combination of reference genes can be found in the literature (Rhinn et al., 2008). For example, Jorgensen and colleagues (2006) reported that 50% of qPCR studies in fish, use the cytoskeletal component *β-actin* (*actb*) as a single reference gene, 30% use the *ribosomal* 18S RNA (18S rRNA) and the remaining 10 % use *elongation factor 1-α* (*ef-1-α*) involved in protein biosynthesis, and *glyceraldehyde-3-phosphate dehydrogenase* (*gapdh*) involved in glucose metabolism. Most of these studies do not provide any evidence on validation (Gutierrez et al., 2008). In addition, nowadays it is widely known that transcription levels of many of these genes vary under different conditions. In fish liver and gonads, for example, *actb* transcription levels are significantly altered after fish exposure to estradiol (Filby and Tyler, 2007) and 18S rRNA and *ef-1-α* do not appear to be appropriate options for the study of dynamic and variable organs such as fish ovaries (Mittelholzer et al., 2007; Deloffre et al., 2012; Diaz de Cerio et al., 2012). In this context, many studies have pointed the advantages of using the amount of cDNA for qPCR data correction (Lundby et al., 2005; Filby and Tyler, 2007; Kroupova et al., 2011; De Santis et al., 2011; Rojo-Bartolomé et al., 2016). In fact, this method is cheaper than the use of reference genes and takes into account possible errors during retrotranscription. Therefore, considering the exact quantity of cDNA used for each qPCR reaction, the variability that could have been generated before qPCRs is negligible (Rhinn et al., 2008).

The use of internal reference genes offers a major advantage because it helps control for technical variation introduced during experimental procedures. Consequently, this strategy has become the most widely adopted approach for normalizing transcriptional data and is explicitly recommended in the MIQE guidelines (Bustin et al., 2025). Despite its widespread use, however, this method presents important limitations. Reference genes must be selected with particular care and must satisfy at least three key criteria: (1) they should exhibit stable transcription across different cell types, (2) remain unaffected by experimental conditions, and (3) display expression levels comparable to those of the target genes (Zhu et al., 2008). Because no single gene consistently meets all these requirements, validation is necessary for each experiment, species, tissue or cell type, and experimental condition (Gutierrez et al., 2008; Huggett et al., 2005). In line with this, the MIQE guidelines discourage the use of unvalidated reference genes and recommend employing multiple reference genes, allowing reliance on a single gene only in clearly justified situations (Bustin et al., 2009; 2025). Although these procedures increase experimental costs (Libus and Štorchová, 2006), their implementation has led to a growing number of studies that adopt combinations of reference genes (Rhinn et al., 2008 Djari et al., 2024; Teixeira et al., 2024). Nevertheless, many qPCR studies continue to rely on single, often unvalidated reference genes. In fish qPCR studies, traditional reference genes such as β-actin (*actb*), *18S rRNA*, and elongation factor 1-α (*ef-1α*) or glyceraldehyde-3-phosphate dehydrogenase (*gapdh*) remain among the most commonly evaluated candidates for normalization; however, their expression stability varies substantially among tissues and experimental conditions, making them unsuitable as universal reference genes (Chen et al., 2025; Wang et al., 2015). Contemporary analyses therefore require systematic validation and often favor combinations of reference genes tailored to each species and context (Wang et al., 2018). Moreover, it is now well established that the transcription of many of these genes varies across physiological and environmental conditions. In fish liver and gonads, for instance, *actb* expression was significantly altered following exposure to ethinyestradiol (Filby and Tyler, 2007), while *18S rRNA* and *ef1α* have proven unsuitable for highly dynamic tissues such as fish ovaries (Mittelholzer et al., 2007; Deloffre et al., 2012; Díaz de Cerio et al., 2012).

Given these limitations, increasing attention has been directed toward normalizing qPCR data using the absolute quantity of cDNA rather than reference genes (Lundby et al., 2005; Filby and Tyler, 2007; Kroupova et al., 2011; De Santis et al., 2011; Rojo-Bartolomé et al., 2016; Nzioka et al., 2023). This alternative is not only more cost-effective but also accounts for variability introduced during the reverse transcription step. By precisely quantifying the amount of cDNA added to each qPCR reaction, variability arising before amplification becomes negligible (Rhinn et al., 2008; De Santis et al., 2011; Rojo-Bartolomé et al., 2016).

The aim of this study is to evaluate four distinct normalization methods for analyzing gene transcription levels in the ovaries of female mullets (*Chelon labrosus*) throughout a complete reproductive cycle. The normalization strategies assessed were: (a) normalization based on the quantified amount of cDNA; (b) normalization using the transcription level of a single commonly employed reference gene (*18S rRNA*, *actb*, *ef-1α*, or *gapdh*); (c) normalization based on the arithmetic mean of the transcription levels of the four reference genes; and (d) normalization using the geometric mean of the transcription levels of all analyzed genes (reference and target genes combined). Three genes associated with oogenesis were selected as targets (Sardi et al., 2015): the acute steroidogenic regulatory protein (*star*), which encodes the protein responsible for translocating cholesterol from the outer to the inner mitochondrial membrane; ovarian aromatase (*cyp19a1a*), which encodes the enzyme involved in estradiol synthesis; and 11β-hydroxylase (*cyp11b1*), which encodes the mitochondrial enzyme required for androgen production. In accordance with the MIQE guidelines, the geNorm and NormFinder statistical tools were employed to validate the stability and homogeneity of transcription for each reference gene, as well as for the geometric mean of the reference genes and the geometric mean of all analyzed genes, thereby ensuring rigor and reliability in the normalization approaches applied.

## MATERIALS AND METHODS

### Source of ovary samples

This study examined the ovary samples of 43 thicklip grey mullets *Chelon labrosus* obtained from the repositories of the Biscay Bay Environmental Biospecimen Bank (BBEBB, UPV/EHU). Fish were collected between September 2010 and September 2011 from the Pasaia–Trintxerpe mooring area (43°19’35’’ N, 1°55’9’’ E), spanning an entire gametogenic and reproductive cycle. Ovary samples were maintained embedded in RNAlater® (Life Technologies, Carlsbad, USA) and frozen in liquid nitrogen. A portion of the ovary tissue was preserved in parallel in methacrylate resin (Technovit 7100; Heraeus Kulzer GmbH & Co., Werheim, Germany) blocks for histological examination and classification according to the gametogenic stage. Fixed ovary samples were sectioned (5 µm) and stained with hematoxylin / eosin. The ovary phase was determined by light microscopy (Olympus BX61), by adapting the protocol described in Bizarro et al., 2014 and Rojo et al., 2016 (Table 1).

**Table 1:** Characteristics of each female gametogenic phase and the number of samples classified in each phase.

| Stage | Name of the stage | Characteristics | Number of females |
| --- | --- | --- | --- |
| R | Regression | Ovary almost empty. Presence of atretic oocytes. Presence of grouped muscle and connective tissues. Thin ovarian wall. | 6 |
| Pv | Previtellogenesis | Presence of perinuclear oocytes. Absence of muscle and connective tissue reduced. Thin ovarian wall. | 19 |
| Ca | Cortical alveoli | Presence of cortical oocytes. Some oocytes in vitellogenesis (less than 50%) | 9 |
| V | Vitellogenesis | Reserve substances in the cytoplasm. Visible extension of the plasma membrane (more than 50% of the oocytes are in vitellogenesis) | 9 |
| M | Mature | Presence of hydrated oocytes. Visible lipid droplets in the cytoplasm | 0 |
|  |  |  | Total: 43 |

### Extraction of RNA and synthesis of cDNA

Total RNA was extracted from the frozen ovarian samples using TRIzol® reagent (Life Technologies, Thermo Fisher Scientific, California, USA) following the manufacturer’s protocol. The resulting RNA was resuspended in 90 µL of RNase / DNase-free water. To ensure high RNA quality, the extracts were treated with DNase I (RNase-Free DNase Set, Qiagen, California, USA) and subsequently purified using the RNeasy® Kit (Qiagen). RNA concentration and purity were assessed spectrophotometrically (Biophotometer, Eppendorf), and only samples displaying absorbance ratios of 260/280 nm between 1.80 and 2.00 and 260/230 nm between 2.00 and 2.20 were retained for downstream analyses. cDNA synthesis was then performed using 1 µg of total RNA per reaction, following the manufacturer’s instructions for the SuperScript® First-Strand Synthesis System (Life Technologies).

The quantity of cDNA was determined fluorometrically using the Quant-iT™ OliGreen® ssDNA Assay Kit (Life Technologies). Fluorescence measurements were performed in a Synergy HT Multi-Mode Microplate Reader (BioTek, Winooski, USA) using 96-well plates (Corning Incorporated, New York, USA). The high-range standard curve supplied with the kit was employed for quantification. Theoretical cDNA concentrations ranging from 0.02 to 0.2 ng/µL were used in a total reaction volume of 100 µL. Fluorescence was recorded at an excitation wavelength of 485/2 nm and an emission wavelength of 528/20 nm.

### Quantification of gene transcription levels

Specific primers for the reference genes *18S rRNA* (AY836368), *actb* (AY836369), *ef-1α* (AY825252), and *gapdh* (KX132908), and the target genes *star* (JX294414), *cyp11b1* (JX294416), and *cyp19a1a* (EF535845) were designed for qPCR analyses (Table 2). Gene transcription levels were quantified in 96-well plates (Applied Biosystems, Thermo Fisher Scientific) using a 7300 Real-Time PCR System thermocycler (Applied Biosystems). Each sample was analysed in triplicate, with reactions containing 10 µL of SYBR® Green PCR Master Mix (Roche, Basel, Switzerland) and 2 µL of cDNA, in a final qPCR volume of 18 µL. Primers were diluted in RNase / DNase-free water and added at the specific concentrations indicated in Table 2. Amplification consisted of 40 cycles under the following conditions: an initial incubation at 50 °C for 2 min, followed by 10 min at 95 °C, and then 40 cycles of denaturation at 95 °C for 15 s and annealing for 1 min at the corresponding primer-specific temperature. A dissociation step was added at the end (95°C for 15 sec), followed by an anneling step for a minute at each gene annealing temperature. Finally, samples were incubated at 95°C for 15 sec.

**Table 2:**
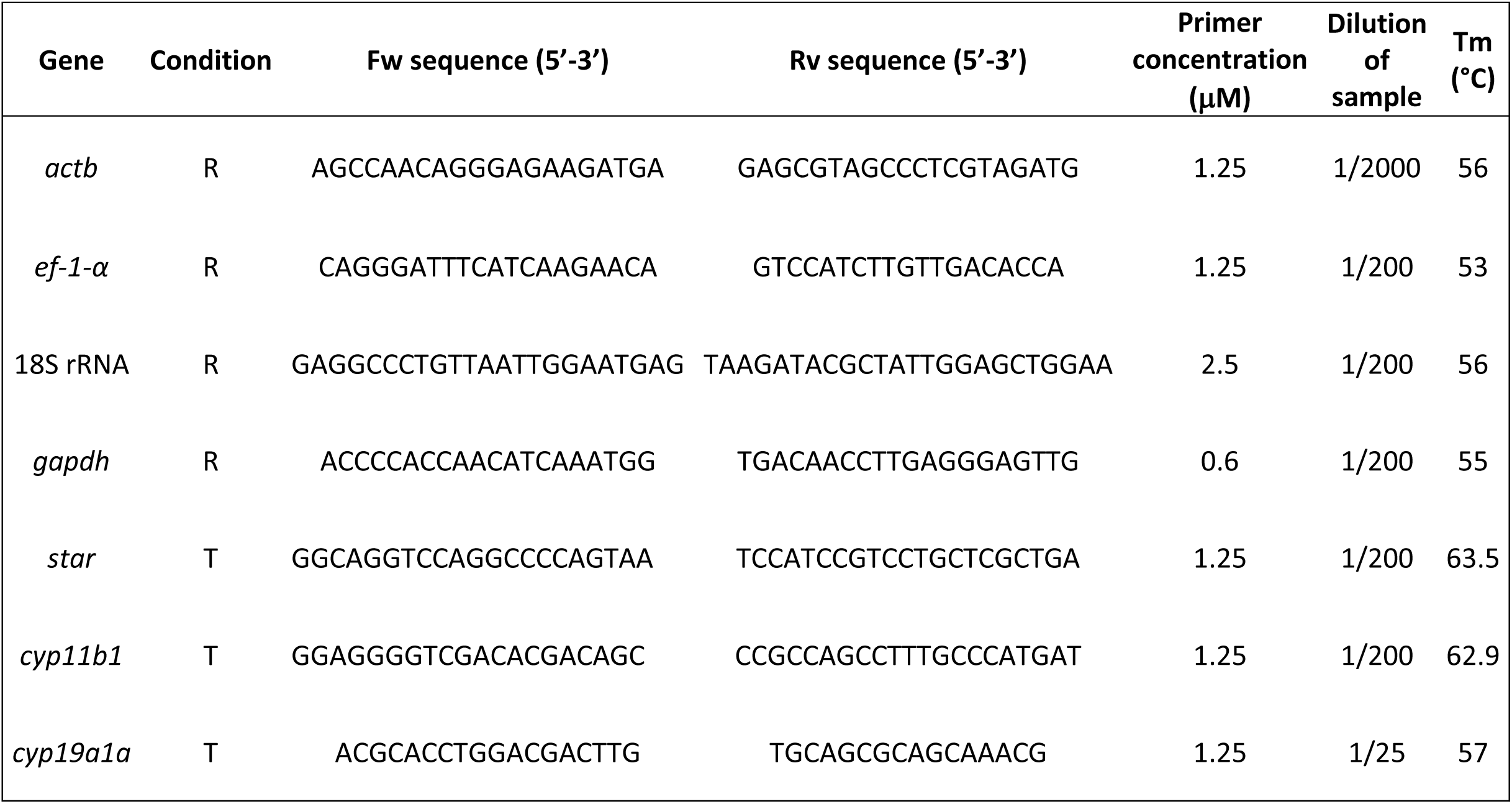
Characteristics of primers and qPCR conditions. Assigned condition to each gene; R: denotes reference genes and T: target genes. Fw: forward, and Rv: reverse primers. (° C): Corresponds to the annealing temperature (Tm: Temperature).

Relative gene expression levels were calculated using the ΔΔCT method (Pfaffl et al., 2001) and the adapted ΔCT procedure described by Rojo-Bartolomé et al. (2016). Reference gene CT values were corrected according to cDNA concentration and cross-normalised against the remaining reference genes, while target genes were normalised to cDNA concentration and each individual reference gene, as well as to the geometric mean of all reference genes (Eg) and the overall mean expression of all genes analysed (EIg). Gene expression results were subsequently grouped according to gametogenic phase (see Table 1); however, no females in the M phase (mature) were obtained.

### cDNA-based correction and efficiency adjustment of gene expression data

Correction of gene transcription levels based on cDNA quantity was performed following the adaptation of the ΔCT method described by Rojo-Bartolomé et al. (2016). For this purpose, the following equation was applied, incorporating the total amount of cDNA loaded into each qPCR reaction:

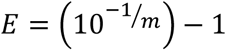

*E* correction of the efficiency and *m* slope of the standard curve.

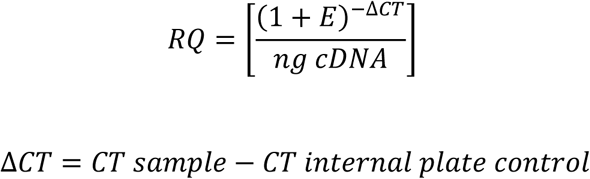

The quantified cDNA content was used both to assess the stability of the candidate reference genes and to correct the transcription levels of all analysed genes. The mean value of a pool of all samples was used as internal plate control.

### Computational analysis of reference gene stability using geNorm and NormFinder

The variability of the candidate reference genes was evaluated using geNorm (Vandesompele et al., 2002; https://genorm.cmgg.be/) and NormFinder (Andersen et al., 2004; http://moma.dk/normfinder-software). Each algorithm applies a distinct statistical approach to determine the most suitable reference gene or gene combination for normalization. geNorm ranks genes according to their transcriptional stability by performing pairwise comparisons and calculating a stability coefficient (M value), with M < 1.5 considered the threshold for an acceptable reference gene; lower M values indicate higher stability. NormFinder provides an ordered list of genes based on their stability and additionally identifies the most stable pairwise combination of reference genes. Data analysis with both tools was carried out strictly following the procedures recommended by their respective authors.

### Statistical analysis

The data were analysed using SPSS v.29 (IBM Corp.). Statistical differences among groups were assessed using the Kruskal–Wallis test, followed by Dunn’s post hoc test for pairwise comparisons. Significance was accepted at p < 0.05. Results were represented using box-and-whisker plots, in which the boxes represent the interquartile range (25th–75th percentiles), the central line denotes the median, and the whiskers indicate the minimum and maximum values.

## RESULTS

### Reference gene stability analysis by geNorm and NormFinder

Overall, none of the selected reference genes met the geNorm stability threshold of M < 1.5. However, most candidates exhibited M values within a narrow range (1.510–1.581), following the order: Eg < *actb* < *gapdh* < *ef-1α* < *EIg* < *18S rRNA* (Section A, Table 3). Among these, *18S rRNA* showed the greatest instability, with an M value of 1.814. The geometric mean of all reference genes (*Eg*) produced an M value of 1.495, falling just below the geNorm limit. In contrast, when target genes were incorporated into the analysis (*EIg*), the M value increased to 1.755, thereby exceeding the acceptable threshold (Table 3). Reference gene stability was further evaluated using NormFinder. According to this algorithm, the ranking of transcriptional stability was: *ef-1α* < *actb* < *Eg* < *gapdh* < *EIg* < *18S rRNA* (Section B, Table 3). The most stable gene pair identified by NormFinder was *actb* and *ef-1α*, with a stability value of 4.793. Consistent with the geNorm analysis, *18S rRNA* was again identified as the least stable reference gene.

**Table 3.** Stability results obtained using geNorm (A) and NormFinder (B) tools. M: stability coefficient calculated by geNorm. Stability: stability value calculated by NormFinder. Eg: geometric mean of reference genes. EIg: geometric mean of all genes, including target genes. The value highlighted in bold corresponds to the gene with the most stable transcription.

| geNorm | 18S rRNA | *actb* | *ef-1-α* | *gapdh* | Eg | Elg |
| --- | --- | --- | --- | --- | --- | --- |
| M<1.5 | 1.814 | 1.510 | 1.581 | 1.552 | **1.495** | 1.755 |

B)
| Normfinder | 18S rRNA | *actb* | *ef-1-α* | *gapdh* | Eg | Elg |
| --- | --- | --- | --- | --- | --- | --- |
| Stability | 7.168 | 6.798 | **6.755** | 7.013 | 6.867 | 7.136 |

### Reference gene stability analysis with cDNA normalization

The stability analysis of the reference genes corrected by cDNA quantification revealed consistent transcription levels for *actb*, *ef-1α*, and *gapdh* throughout oogenesis (Fig 1B–D). In contrast, *18S rRNA* exhibited a progressive increase in transcription as oogenesis advanced, with the lowest expression observed during the previtellogenic phase (Pv) and the highest during vitellogenesis (V) (Fig 1A). Accordingly, the ranking of genes from most to least stable was: *actb* < *ef-1α* < *gapdh* < *18S rRNA*. Despite the pronounced variability of *18S rRNA* expression across the oogenic phases, this gene was nevertheless included as a reference in subsequent analyses to evaluate the potential impact that its instability might have on the interpretation of target gene transcription results.

**Fig 1.**
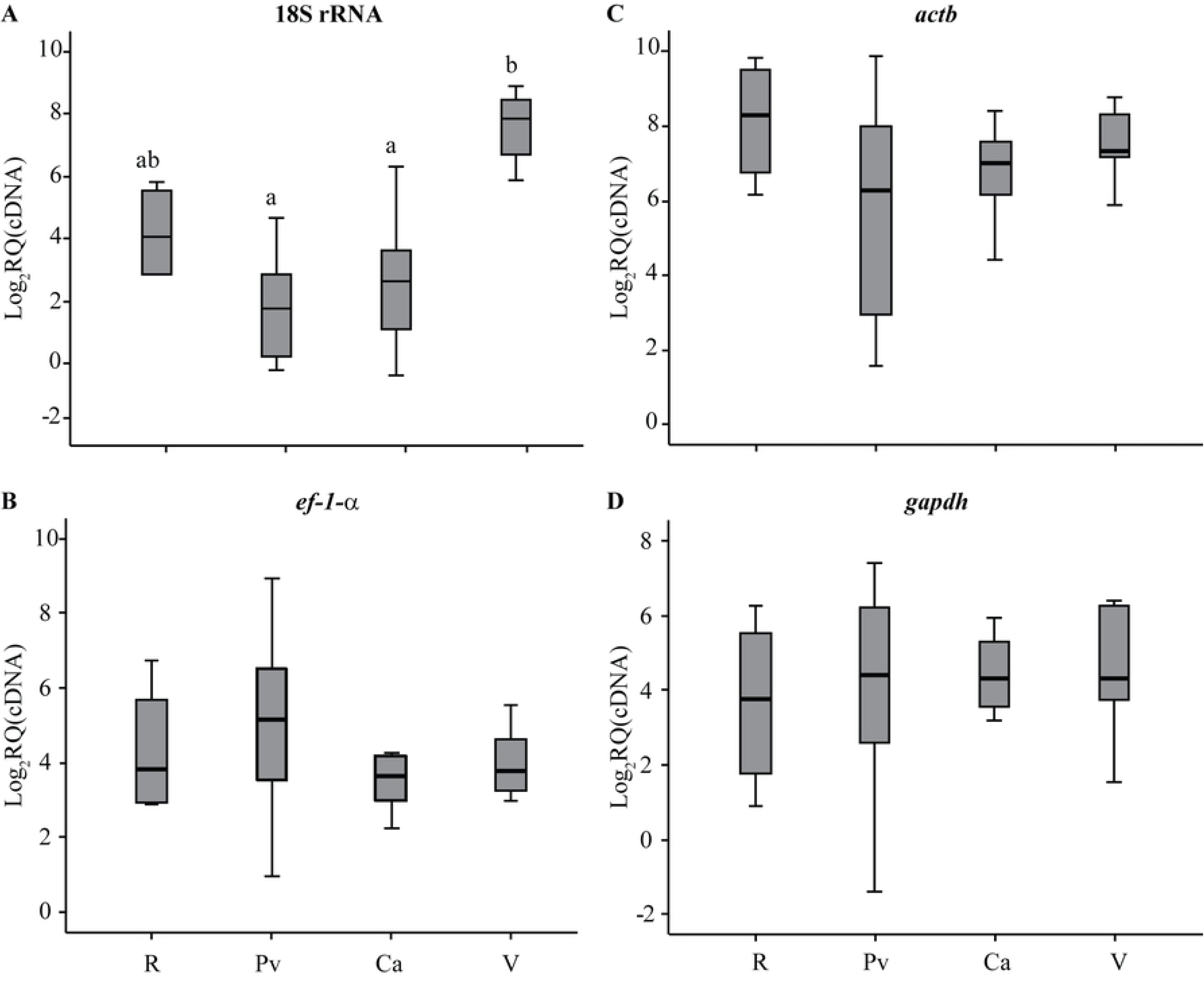
Stability of transcription levels of selected reference genes based on cDNA quantity. Ovarian developmental stages are indicated on the X-axis using the following abbreviations: E = regression; Pb = previtellogenesis; Ak = cortical alveoli; V = vitellogenesis. Log2RQ: Fold Change in relative quantity of each transcript. Panels: A) *18S rRNA*, B) *ef-1-α*, C) *actb*, D) *gapdh*. Different letters indicate statistically significant differences between groups (p < 0.05).

### Transcription levels of reference genes during oogenesis corrected using alternative reference gene candidates

When reference-gene transcription levels were corrected using different reference genes, *18S rRNA* consistently displayed an upward trend throughout oogenesis, irrespective of the gene used for normalisation (Fig 2). The lowest transcription levels were observed during the previtellogenic phase (Pv), whereas the highest levels occurred during vitellogenesis (V). By contrast, the transcription patterns of the remaining reference genes varied depending on the reference gene selected for data normalisation (Figs 3–5). For *actb* (Fig 3), normalisation against *ef-1α* (Fig 3C) resulted in relatively stable transcription across all oogenic phases. However, when *actb* was normalised to *gapdh*, transcription levels appeared elevated during the regression phase (R) compared with the other phases (Fig 3B). Conversely, normalisation against *18S rRNA* produced a decreasing trend in *actb* transcription (Fig 3A), with the highest levels during previtellogenesis and the lowest during vitellogenesis, mirroring the low and high *18S rRNA* transcription levels shown in Fig 2. The transcription profile of *ef-1α* (Fig 4) followed a similar pattern to that of *actb* when normalised against *ef-1α*-independent stable genes (Fig 3C). Normalisation against *gapdh* produced an upward trend across oogenesis, with the highest *ef-1α* transcription levels observed during vitellogenesis (Fig 4B). When normalised against *18S rRNA*, *ef-1α* exhibited a downward trend as oogenesis progressed (Fig 4A), again resembling the pattern observed for *actb* (Fig 3A). For *gapdh* (Fig 5), normalisation using *ef-1α* or *actb* (Fig 5B–C) revealed largely stable transcription throughout oogenesis, although a slight decrease was detected during the regression phase (R). In contrast, normalisation against *18S rRNA* resulted in a marked downward trend in *gapdh* transcription (Fig 5A), with the lowest levels occurring during vitellogenesis, coinciding with the highest *18S rRNA* transcription levels (Fig 2).

**Fig 2.**
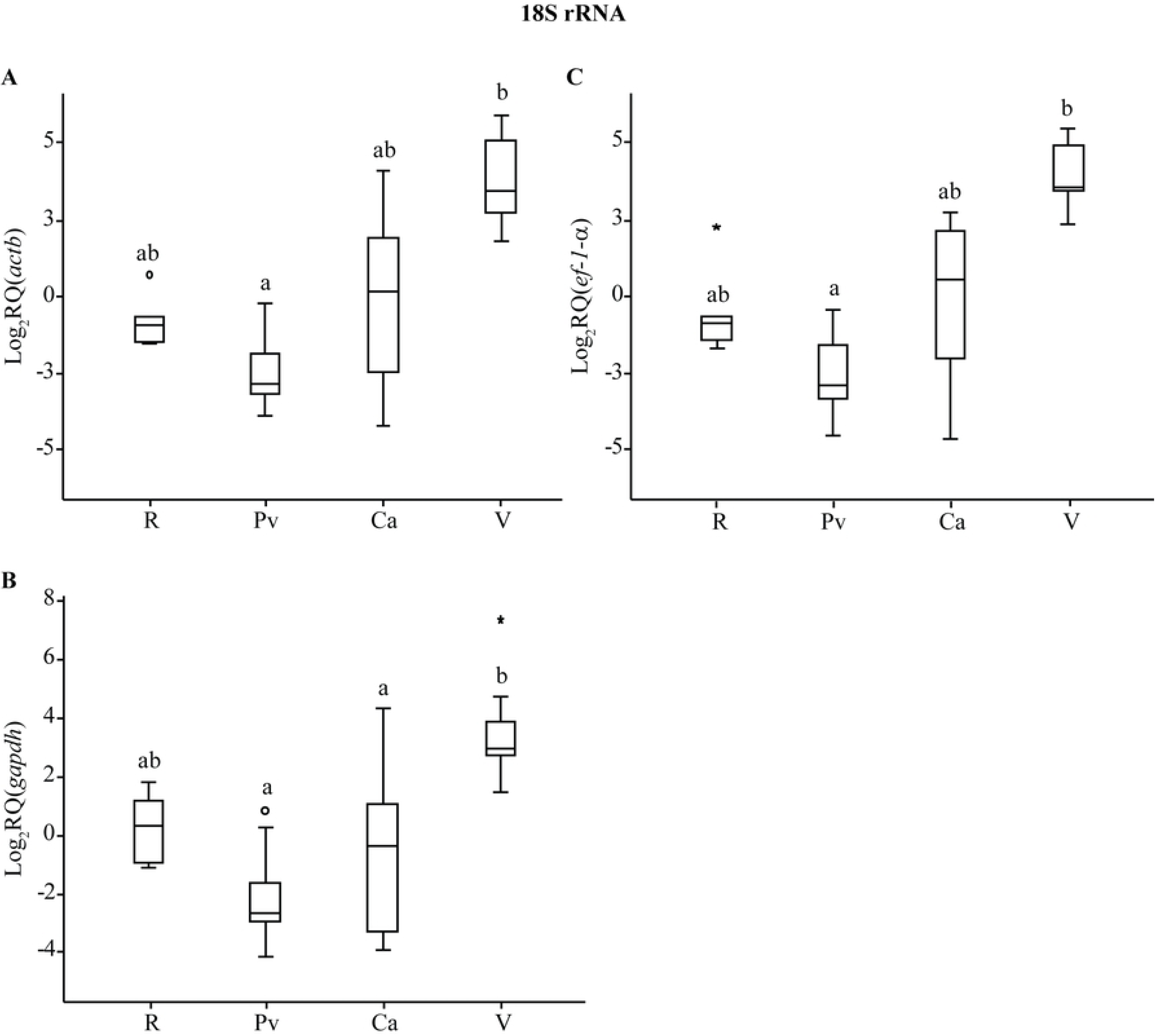
Relative transcription levels of *18S rRNA* normalised using different reference genes. Ovarian developmental stages are indicated on the X-axis using the following abbreviations. Log2RQ: Fold Change in relative quantity of each transcript. The reference gene used for normalization is shown in parentheses: A) *actb*, B) *gapdh*, and C) *ef-1-α*. Different letters indicate statistically significant differences between groups (p < 0.05).

**Fig 3.**
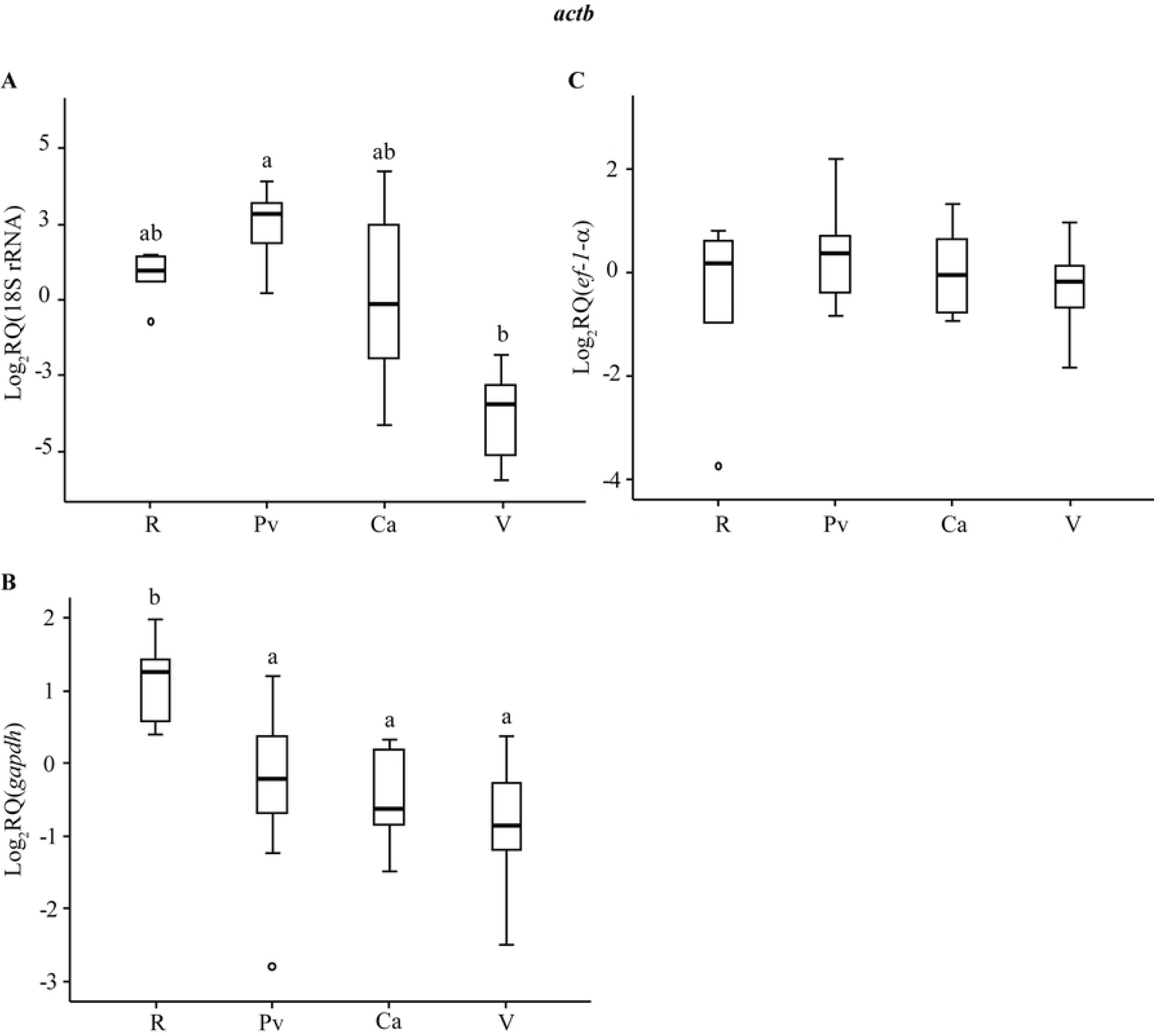
Relative transcription levels of *actb* normalised using different reference genes. Ovarian developmental stages are indicated on the X-axis using letter codes. Log2RQ: Fold Change in relative quantity of each transcript. The reference gene used for normalization is shown in parentheses: A) *18S rRNA*, B) *gapdh*, and C) *ef-1-α*. Different letters indicate statistically significant differences between groups (p < 0.05).

**Fig 4.**
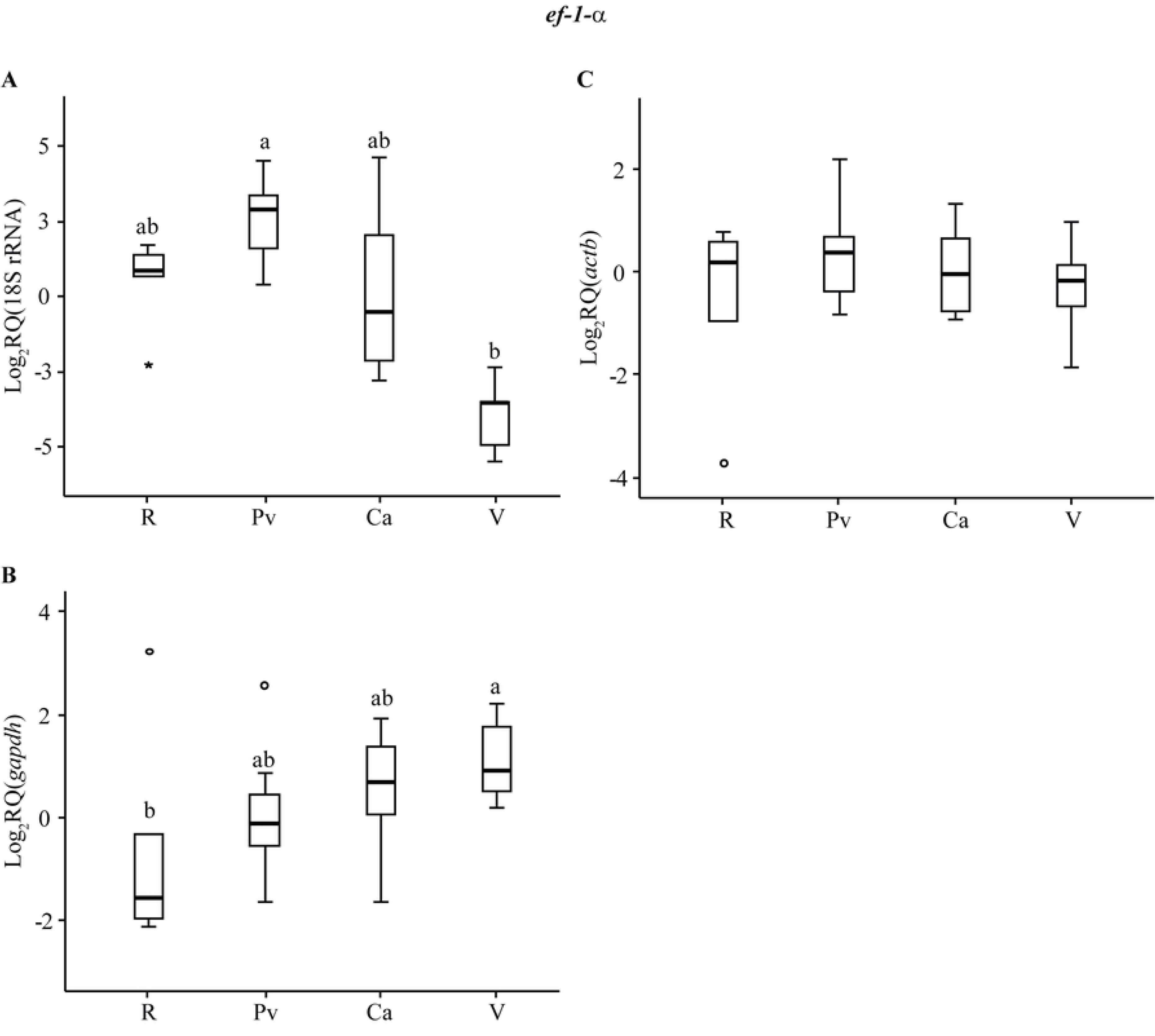
Relative transcription levels of *ef-1-α* normalised using different reference genes. Ovarian developmental stages are indicated on the X-axis using letter codes. Log2RQ: Fold Change in relative quantity of each transcript. The reference gene used for normalization is shown in parentheses: A) *18S rRNA*, B) *gapdh*, and C) *actb*. Different letters indicate statistically significant differences between groups (p < 0.05).

**Fig 5.**
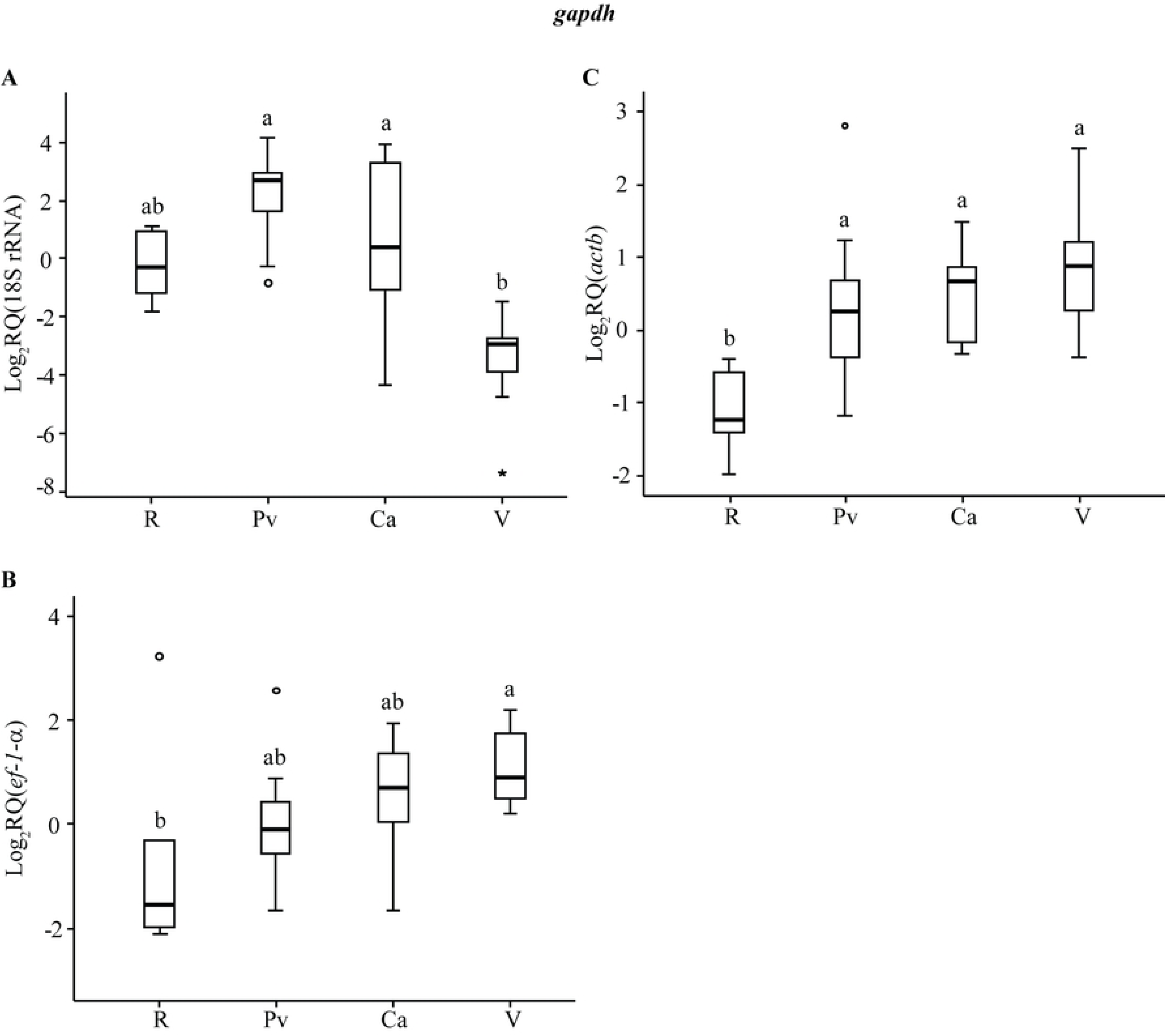
Relative transcription levels of *gapdh* normalised using different reference genes. Ovarian developmental stages are indicated on the X-axis using letter codes. Log2RQ: Fold Change in relative quantity of each transcript. The reference gene used for normalization is shown in parentheses: A) *18S rRNA*, B) *ef-1-α*, and C) *actb*. Different letters indicate statistically significant differences between groups (p < 0.05).

### Comparison of cDNA-based, reference gene–based, and geometric mean–based normalisation of target gene transcription during oogenesis

*star* transcription displayed a progressive increase throughout oogenesis, reaching its highest levels during the vitellogenic phase (V) when normalised to cDNA quantity or when using *actb* and *ef-1α* as reference genes (Fig 6A, C, D). A similar pattern was observed when *star* was normalised using the geometric mean of all reference genes (Eg) and of all reference and target genes (EIg) (Fig 6F, G). In contrast, normalisation with *gapdh* or *18S rRNA* resulted in no detectable changes in *star* transcription across the oogenic stages (Fig 6B, E).

**Fig 6.**
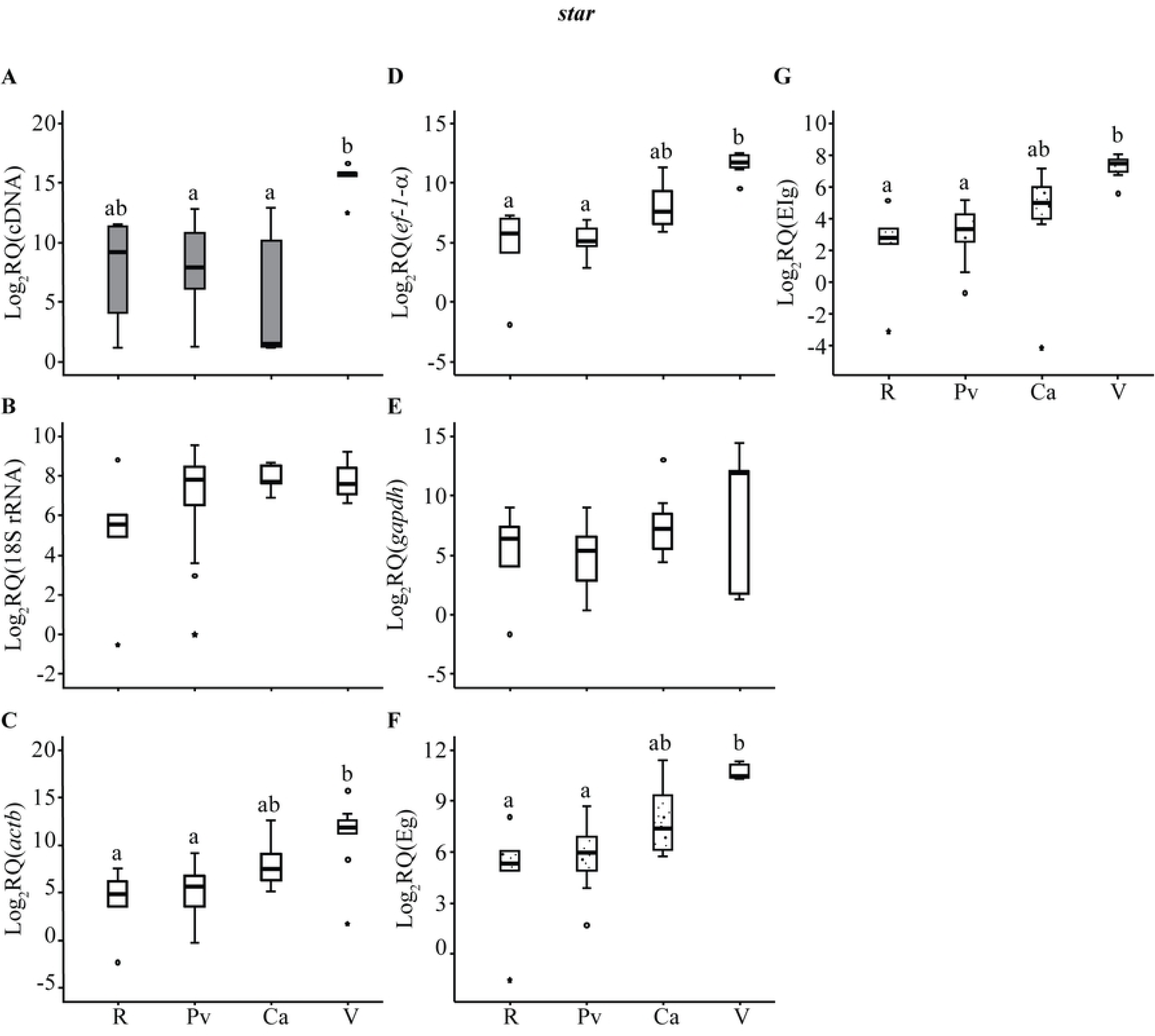
Relative transcription levels of *star* normalised using cDNA and different reference genes. Ovarian developmental stages are indicated on the X-axis using letter codes. Log2RQ: Fold Change in relative quantity of each transcript. The normalization criterion used is shown in parentheses: A) cDNA, B) *18S rRNA*, C) *actb*, D) *ef-1-α*, E) *gapdh*, F) Eg (geometric mean of reference genes), and G) EIg (geometric mean of all genes). Different letters indicate statistically significant differences between groups (p < 0.05).

For *cyp11b1*, no statistically significant variations were observed across oogenesis when normalised to cDNA quantity (Fig 7A). Normalisation using individual reference genes, as well as Eg and EIg, generally yielded stable transcription levels throughout the cycle, although some significant pairwise differences were detected. Overall, all reference-gene–based normalisation methods—except normalisation with *18S rRNA* (Fig 7B)—indicated that *cyp11b1* transcription remained consistent between early (Pv) and late oogenic (V) phases. Notably, the regression phase (R) showed significantly elevated transcription, whereas the cortical alveoli phase (Ca) exhibited the lowest levels.

**Fig 7.**
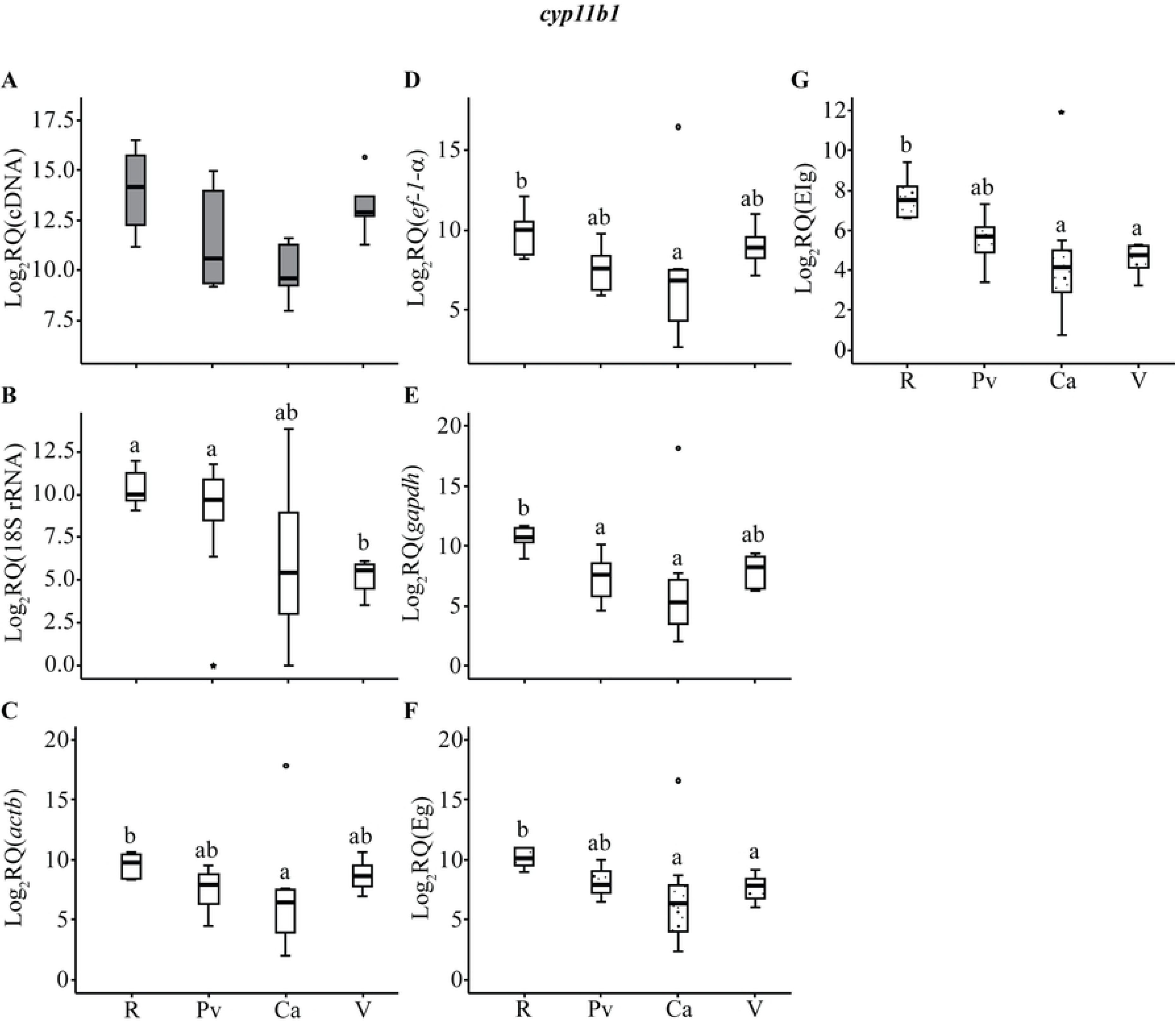
Relative transcription levels of *cyp11b1* normalised using cDNA and different reference genes. Ovarian developmental stages are indicated on the X-axis using letter codes. Log2RQ: Fold Change in relative quantity of each transcript. The normalization criterion used is shown in parentheses: A) cDNA, B) *18S rRNA*, C) *actb*, D) *ef-1-α*, E) *gapdh*, F) Eg (geometric mean of reference genes), and G) EIg (geometric mean of all genes). Different letters indicate statistically significant differences between groups (p < 0.05).

*cyp19a1a* transcription increased progressively during oogenesis, with the highest expression recorded during the cortical alveoli phase, regardless of whether the data were normalised to cDNA quantity or to the stable reference genes (Fig 8A, C–G). As with the other genes, normalisation with *18S rRNA* masked these transcriptional changes due to the opposing expression pattern of *cyp19a1a* and *18S rRNA* (Fig 8B).

**Fig 8.**
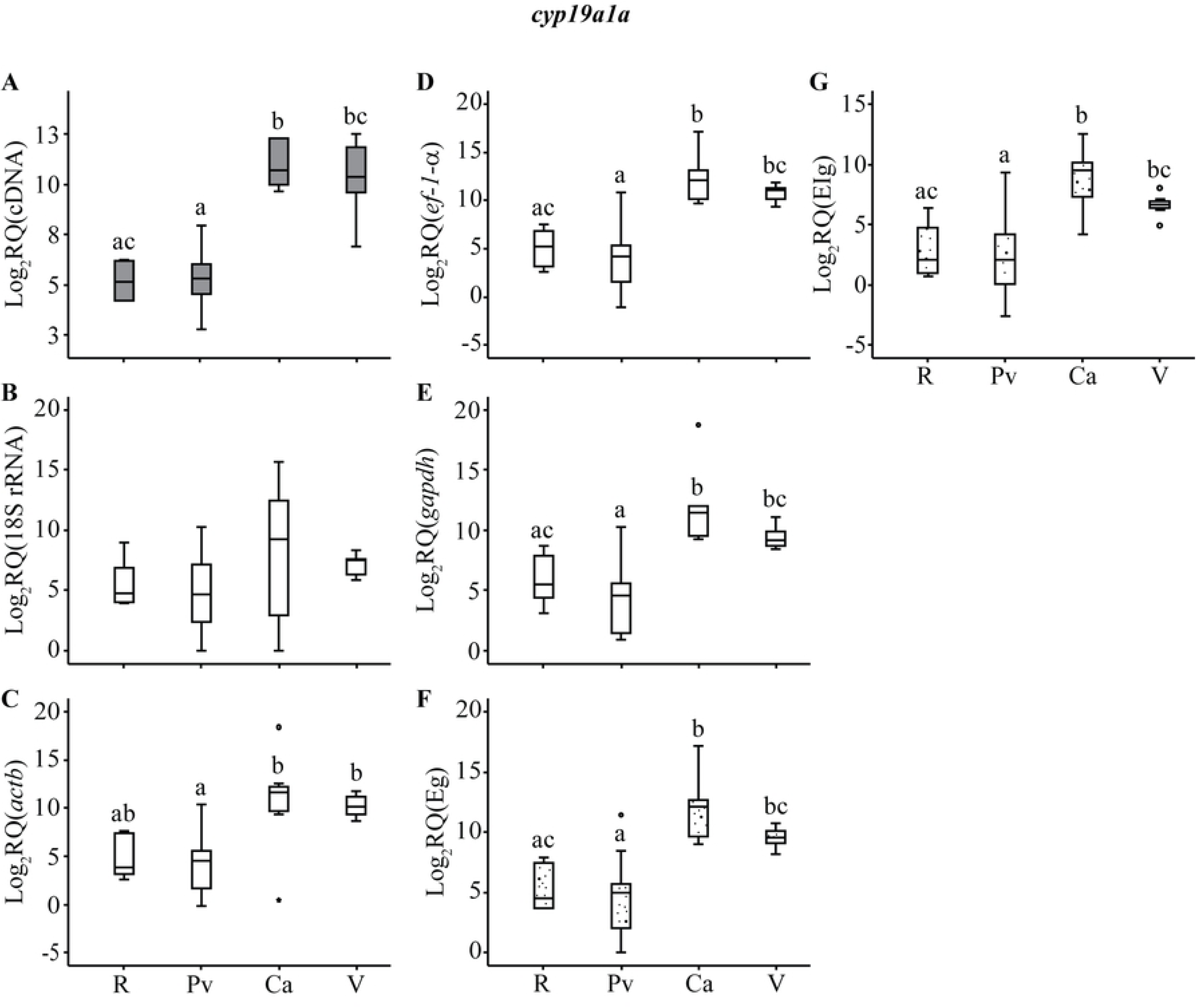
Relative transcription levels of *cyp19a1a* normalised using cDNA and different reference genes. Ovarian developmental stages are indicated on the X-axis using letter codes. Log2RQ: Fold Change in relative quantity of each transcript. The normalization criterion used is shown in parentheses: A) cDNA, B) *18S rRNA*, C) *actb*, D) *ef-1-α*, E) *gapdh*, F) Eg (geometric mean of reference genes), and G) EIg (geometric mean of all genes). Different letters indicate statistically significant differences between groups (p < 0.05).

Across all three target genes analysed, normalisation using the geometric mean approaches (Eg and EIg) produced transcription profiles closely matching those obtained using cDNA quantity. Furthermore, each target gene exhibited the same expression trend when normalised with either of the geometric-mean-based methods (Fig 6F–G; Fig 7F–G; Fig 8F–G).

## DISCUSSION

Reliable normalization is essential for accurate RT-qPCR quantification, and current standards emphasize empirical validation of any chosen strategy. Our results demonstrate that reference-gene stability during *Chelon labrosus* oogenesis is strongly stage-dependent, reflecting the extensive transcriptional and cellular remodeling characteristic of teleost ovarian development. This variability confirms that no reference gene can be assumed stable across oogenic stages and underscores the need for design specific evaluation.

Across the methods used to evaluate reference gene transcription stability, *actb* and *ef-1α* emerged as the most stable candidates, while *18S rRNA* consistently showed poor performance. The instability of *18S rRNA* is mechanistically consistent with dynamic ribosome biogenesis during oogenesis, indicating that rRNA-based normalization is inappropriate for this biological process (Kroupova et al., 2011; Ortiz-Zarragoitia et al., 2014; Rojo-Bartolomé et al., 2016 and 2017; Locati et al., 2017). *gapdh*, although stable in earlier stages, exhibited clear variability during regression and is therefore unsuitable for full-cycle analyses.

Normalization to measured cDNA quantity produced expression patterns that closely matched those derived from validated reference genes, suggesting that cDNA-based approaches can capture the underlying biological dynamics when appropriate quality controls are implemented. This offers a practical alternative in systems where reference gene validation is challenging or when resource efficiency is a priority (Filby and Tyler, 2007; Rojo-Bartolomé et al., 2016 and 2017). Nevertheless, the approach has inherent limitations—particularly sensitivity to RNA integrity and reverse-transcription variability—which must be addressed through rigorous experimental control.

The transcriptional patterns of *star*, *cyp19a1a*, and *cyp11b1* corroborate their established roles in teleost ovarian physiology. The consistent upregulation of *star* and *cyp19a1a* during vitellogenic progression aligns with their functions in steroidogenic activation and estradiol synthesis, respectively (Nakamura et al., 2005; Villeneuve et al., 2010; Sardi et al., 2015). In contrast, *cyp11b1* remained low throughout oogenesis, with only a modest increase during regression, reflecting its predominantly testis-biased expression and limited involvement in ovarian steroidogenesis (Blázquez et al., 2009; Zhang et al., 2010; Sardi et al., 2015).

Comparing normalization approaches highlights several trade-offs. Multi-gene geometric means provide robust correction but require substantial investment in gene validation and increase analytical complexity, particularly when few target genes are examined (Libus & Štorchová, 2006; Hruz et al., 2011; Ling and Salvaterra, 2011). Reference-gene normalization remains effective but labor-intensive and species-dependent (Von Schalburg et al., 2005; Filby and Tyler, 2007; Luckenbach et al., 2008a, b; Tingaud-Sequeira et al., 2009; Gao et al., 2020; Pagano et al., 2024). cDNA-based normalization is simple and cost-efficient, but it requires rigorous quality control and may be less appropriate in contexts with large shifts in RNA composition, such as late oogenesis (Bir et al., 2023). Its accuracy can also be compromised by genomic DNA contamination or RNA degradation. However, these limitations can be minimized through standard procedures, including DNase treatment, careful assessment of RNA integrity, and optimized reverse-transcription workflows.

Overall, these results provide a refined framework for gene expression normalization in teleost oogenesis and identify *actb*, *ef-1α*, cDNA concentration, the geometric mean of reference genes, and the geometric mean of all genes (reference genes plus target genes) suitable approaches for normalising transcription levels during fish oogenesis. More broadly, they highlight the need for normalization strategies that accommodate dynamic developmental contexts. Identifying appropriate reference genes is further complicated in different fish species, in contaminant-exposure experiments, and particularly when analysing transcription of genes associated with development or oogenesis (Von Schalburg et al., 2005; Filby and Tyler, 2007; Luckenbach et al., 2008a, b; Tingaud-Sequeira et al., 2009; Pomianowski and Burzynski 2024; Rosetto Vidal et al., 2024), as sequence information may be unavailable for the species under study (Mittelholzer et al., 2007). The use of the geometric mean of target-gene transcription levels is a suitable method as an alternative normalisation strategy (Heckmann et al., 2011). However, it requires quantifying the transcription levels of several genes to obtain robust results, which could be a limitation when only a few target genes are analysed. In contrast, normalisation to cDNA quantity is straightforward, inexpensive, avoids procedural biases during experimentation, yields biologically coherent results, and requires minimal resources (Libus & Štorchová, 2006; Filby and Tyler, 2007; Rhinn et al., 2008; De Santis et al., 2012; Rojo-Bartolomé et al., 2016, 2017). Future work should prioritize developing standardized, species-independent calibration frameworks tailored to oogenesis that integrate cDNA-based normalization with adaptive reference gene selection algorithms to improve accuracy, reduce labour requirements, and enhance applicability across diverse teleost models.

## ACKNOWLEDGMENTS

This study was funded by the Spanish Ministry of Science, Innovation and Universities (grant PID2023–146085NB-I00) and the Basque Government (CRG grant IT-1743–22). We are thankful to the Biscay Bay Environmental Biospecimen Bank (BBEBB; PiE-UPV/EHU) for providing mullet ovary tissues.

